# Modeling the role of ATP metabolism in articular cartilage and osteoarthritis

**DOI:** 10.1101/2025.07.17.665362

**Authors:** Dhruba Jyoti Mech, Mohd Suhail Rizvi

**Affiliations:** Department of Biomedical Engineering, Indian Institute of Technology Hyderabad, Kandi, Sangareddy, Telangana, 502284, India

**Keywords:** articular cartilage, osteoarthritis, ATP metabolism, HIF-1α

## Abstract

Osteoarthritis, a prevalent degenerative joint disease, is characterized by progressive degradation of articular cartilage. The avascular nature of articular cartilage makes it vulnerable to metabolic disruptions under hypoxic conditions. Central to this process is the role of ATP metabolism in chondrocytes, which generally maintains a delicate balance between glycolysis and oxidative phosphorylation. To investigate the balance between these two mechanisms and their regulation, we developed a comprehensive mathematical model simulating ATP metabolism in chondrocytes. The model incorporates key metabolic regulators, capturing the bistable switching between glycolysis and oxidative phosphorylation under varying nutrient conditions. Our simulation also accounts for stochastic fluctuations in oxygen and glucose levels, mimicking physiological conditions during mechanical loading, and their impact on articular cartilage dynamics. The results demonstrate that chronic hypoxia induces an irreversible metabolic shift to glycolysis, leading to sustained reductions in ATP levels and progressive ECM loss. Interestingly, the model predicts that physiological stochasticity in oxygen levels, representative of mechanical loading during physical activity, enhances metabolic flexibility and promotes ATP synthesis. When testing therapeutic interventions, we found that while exogenous ECM supplementation provides transient matrix restoration, only approaches targeting metabolic dysfunction - either through enhanced ATP synthesis or controlled suppression of regulatory factors - successfully reverse the pathological glycolytic shift. Our model suggests that optimal therapeutic approaches should combine ATP metabolic modulation with structural support to maintain beneficial nutrient fluctuations. The framework provides a basis for the development of personalized treatment strategies that address both the metabolic and structural aspects of osteoarthritis, offering new possibilities for restoring cartilage homeostasis and preventing disease progression.

## 1 Introduction

Osteoarthritis (OA), the most prevalent degenerative joint disease, affects over 240 million people worldwide and is a leading cause of adult disability (Allen et al., 2022). Its hallmark is the progressive breakdown of articular cartilage – the smooth, load-bearing tissue that cushions joints – accompanied by subchondral bone remodeling and synovial inflammation (Hunter and Bierma-Zeinstra) (Fig. 1A). Prevalence of OA rises sharply with age, as approximately 10–12% of adults over 60 experience symptomatic OA, reflecting the cumulative toll of mechanical stress and diminished cartilage repair capacity.

**Figure 1.**
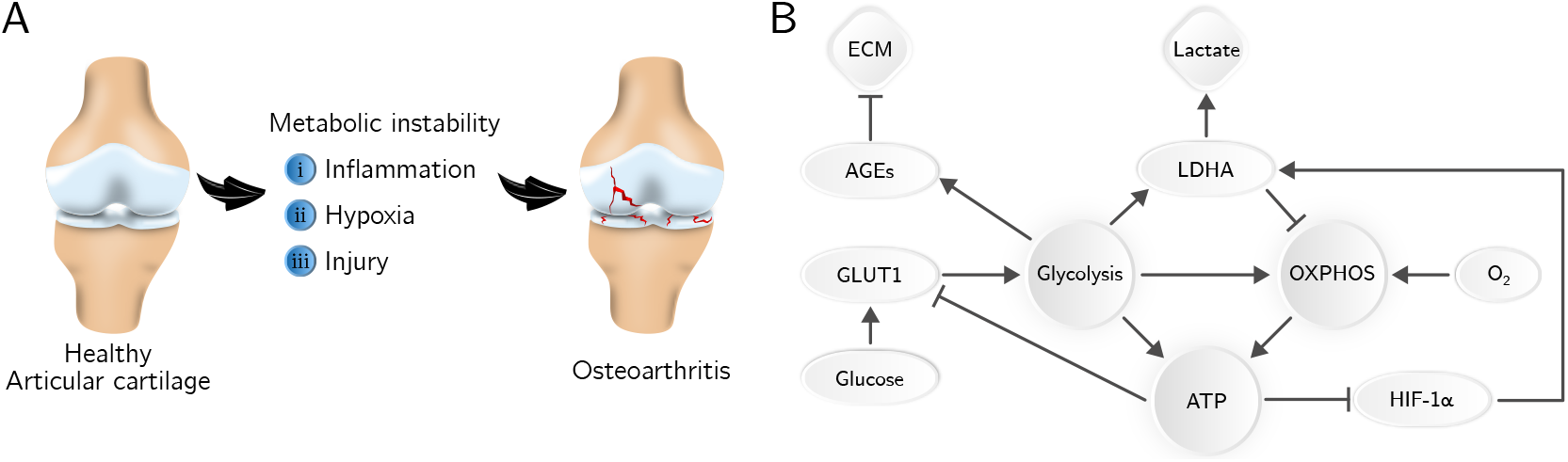
(A) Schematic showing articular cartilage and development of osteoarthritis by instability in ATP metabolism. (B) Regulatory network of ATP metabolism in articular cartilage.

Articular cartilage (AC), a porous connective tissue, provides lubrication (Jahn et al., 2016) and supports load transmission at the articular joints (Kelly and O’Connor, 1996a,b). This avascular tissue is composed of a dense extracellular matrix (ECM) populated sparsely by chondrocytes (Pueyo Moliner et al., 2025). The ECM in AC is primarily made up of water, collagen, and proteoglycans, along with smaller quantities of non-collagenous proteins and glycoproteins (Sophia Fox et al., 2009). Metabolism, particularly energy production through molecules such as ATP, is a fundamental regulator of cellular function across biological systems (Chen et al., 2023; Liu et al., 2025). Under stress conditions – such as hypoxia in the articular cartilage – mammalian cells often undergo metabolic adaptations to sustain energy homeostasis and ensure survival (Rajpurohit et al., 1996; Thoms et al., 2013; Fermor et al., 2010). Typically, chondrocytes rely predominantly on glycolysis for ATP generation (Fig. 1B), with oxidative phosphorylation (OXPHOS) playing a minor role due to the low oxygen availability in the microenvironment of articular cartilage (Martin et al., 2012; Blanco et al., 2020). This metabolic preference supports ECM stability by balancing anabolic and catabolic processes. However, in osteoarthritis (OA), emerging evidence highlights significant disruptions in ATP metabolism, leading to impaired chondrocyte function and progressive ECM degradation (Mobasheri et al., 2017; Wu et al., 2022; Blanco et al., 2020). Interestingly, in cancer biology, the balance between glycolysis and OXPHOS pathways of ATP synthesis has been shown to play a critical role in disease progression (Yu et al., 2017). These parallels suggest that altered energy metabolism in OA may also involve a more nuanced regulation between glycolysis and OXPHOS. Consequently, targeting metabolic pathways, particularly those governing ATP production, represents a promising therapeutic strategy for OA.

Glucose metabolism in chondrocytes starts with glucose transport in cells through GLUT1 (Fig. 1B) (Mobasheri et al., 2002). GLUT1 expression is tightly regulated, increasing under hypoxia or glucose deprivation and decreasing under high-glucose conditions (Pi et al., 2024; Rosa et al., 2009). A major portion of the glucose entering the cells goes through glycolysis (a process with a net gain of 2 ATP molecules), and its end products can either get converted to lactate (a proxy measure of glycolytic activity) to complete the process of glycolysis or move to mitochondria to participate in OXPHOS (net yield of 36 molecules of ATP per molecule of glucose) (Pi et al., 2024). Articular cartilage being avascular tissue, the synovial fluid in the joint is its primary source of glucose. Its relatively hypoxic environment results in lower energy production by OXPHOS. In a situation of aberrant upregulation of GLUT1, excessive glucose influx results in the formation of advanced glycation end products (AGEs). These AGEs accumulate in the ECM, impairing its structural integrity and promoting inflammation,thereby contributing to cartilage degeneration (Saudek and Kay, 2003).

One of the central players in the regulation of this metabolic pathway is the transcription factor hypoxia-inducible factor 1-*α* or HIF-1*α* (Zhang et al., 2015; Fernández-Torres et al., 2017). Stabilized under low oxygen levels, HIF-1*α* orchestrates the metabolic reprogramming of chondrocytes by promoting anaerobic glycolysis over oxidative phosphorylation (Jiang et al., 2024). It does so by upregulating glycolytic genes and enhancing the expression of GLUT1, further driving glucose uptake. HIF-1*α* also induces the expression of LDHA (lactate dehydrogenase A), an enzyme that catalyzes the conversion of pyruvate to lactate (glycolytic end product) (Cui et al., 2017). This conversion regenerates NAD+, sustaining glycolysis and allowing ATP production under anaerobic conditions (Pi et al., 2024). The ECM, composed of collagen and proteoglycans, is sensitive to metabolic alterations in chondrocytes. Disrupted glucose metabolism, particularly the overactivity of GLUT1 and LDHA, has been known to impair ECM synthesis and accelerate its breakdown (Tan et al., 2022; Jiang et al., 2024). Thus, while glycolysis is essential in healthy cartilage, its dysregulation in OA, marked by aberrant GLUT1, persistent HIF-1*α* signaling, and excessive LDHA activity, creates a pathological feedback loop that undermines cartilage integrity.

This demonstrates that the ATP metabolism in chondrocytes is tightly regulated by several key molecular players, and there is strong evidence now to suggest that a disruption in ATP synthesis can be one potential pathway for the progression of osteoarthritis. Mathematical modeling of biochemical and gene regulatory processes has been one of the major successes in the fields of systems biology and biophysics (Ingalls, 2013; Noble, 2010). Mathematical modeling has also been applied to the study of articular cartilage biology and mechanics (Carter and Wong, 2003). In particular, the impact of inflammatory cytokines on chondrocyte physiology and their contribution to the development of osteoarthritis has been explored (Graham et al., 2012). However, most previous studies have primarily focused on the effects of mechanical loading and injury on cartilage physiology and osteoarthritis progression (Wang et al., 2014; Kapitanov et al., 2017). In contrast, the role of glucose metabolism and ATP production in articular cartilage has not been systematically investigated through mathematical modeling.

In this work, we develop a mathematical model to describe glucose metabolism and ATP synthesis in articular cartilage. We demonstrate that ATP production via glucose metabolism is governed by a bistable regulatory switch. Using this framework, we investigate how disruptions in ATP synthesis can lead to reductions in ECM density under conditions of chronic inflammation or mechanical injury. We show that these pathological conditions can trigger a switch in the metabolic state of the system, driving it from a healthy to a diseased stable state and thereby contributing to the onset and progression of osteoarthritis.

## 2 Mathematical model

### 2.1 Basic assumptions

In the mathematical model, we have made several simplifying assumptions to capture the essential dynamics of glucose metabolism in articular cartilage. The tissue is treated as a well-mixed system, neglecting spatial heterogeneity to focus on bulk metabolic interactions. To maintain analytical tractability and focus on core metabolic dynamics, we have ignored spatial heterogeneity within articular cartilage. In particular, depth-dependent gradients of oxygen, glucose, and ECM properties are not taken into account. In this work, we are focusing only on a small subset of proteins involved in the ATP synthesis and its regulation. We will see that despite this simplification, we are able to reproduce several of key experimental observations. The model operates on two distinct timescales: a fast timescale for metabolic reactions and a slower one for genetic regulations. We have also explored the model by varying the parameters in a wide range of values to see that the results reported here are robust. In both metabolic and gene regulatory interactions, all the dynamics are modeled using Hill kinetics, allowing for sigmoidal response curves that approximate biological regulation.

### 2.2 Model formulation

The model is formulated based on different metabolic and gene regulatory interactions that are summarized in Fig. 1B. Since GLUT1 is responsible for the transport of glucose from the extracellular space to the inside, we consider the transport of glucose from outside to inside the cell to be dependent on GLUT1 levels and write

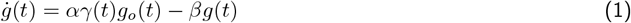

where *γ*(*t*) represents the GLUT1 levels in chondrocytes, and *g*_*o*_(*t*) is the extracellular glucose concentration. Please note that in this equation, we have mentioned the time dependencies. However, in the subsequent equations we will not write it explicitly, even though it is assumed for all metabolites and proteins.

The glucose entering the cell undergoes glycolysis, and some fraction of it also undergoes OXPHOS. Both of these processes produce ATP. The amount of glucose entering OXPHOS depends on the levels of LDHA, the enzyme catalyzing the pyruvate to lactate conversion. Therefore, we write the rate of change in ATP levels as

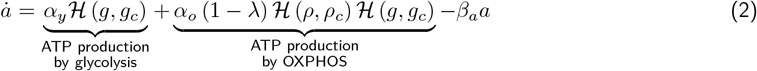

where *ρ* is the available oxygen, *λ* is the level of LDHA, and

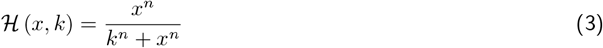

is the Hills function. Please note that for very high levels of LDHA, that is *λ* → 1, all of the glucose is converted to lactate at the end of glycolysis, and there is no OXPHOS.

We also consider the dynamics of GLUT1 and hypothesize that its regulation is governed by the ATP synthesis. This assumption is based on the observations of an increase in the levels of GLUT1 during hypoxia as well as in low glucose levels (Pi et al., 2024; Jiang et al., 2024). We combine both of these mechanisms by considering GLUT1 regulation by ATP alone. This gives

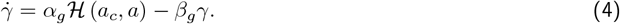

Similar to ATP dependent regulation of GLUT1, we also consider a similar regulation of HIF-1*α* that is known to be stabilized at low ATP levels (Valvona et al., 2016). Therefore, we get

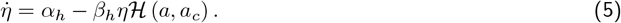

HIF-1*α*, in turn, regulates the levels of LDHA (Valvona et al., 2016), and gives us

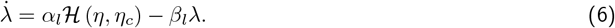

These equations complete the description of the regulation of ATP synthesis in glucose metabolism. To integrate this with cartilage homeostasis, we couple these metabolic dynamics with extracellular matrix remodeling processes. A key link arises from advanced glycation end-products (AGEs), which are produced during glycolytic activity. These AGEs are biologically significant as they directly contribute to ECM degradation (Saudek and Kay, 2003). By incorporating AGE-mediated ECM damage into the framework, the model captures the interplay between metabolism and ECM stability as

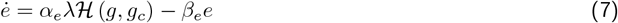

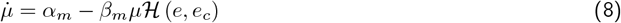

where *e* and *µ* represent AGEs and ECM density, respectively. These equations describe the ATP metabolism and its role in ECM stability in articular cartilage. In the following, we explore the properties of this system under normal and pathological conditions.

### 2.3 Non-dimensionalization and system parameters

We non-dimensionalize concentrations of glucose *g*, oxygen *ρ*, ATP *a*, LDHA *λ*, AGEs *e*, and ECM *µ* by their critical concentrations specified in respective Hill’s function, that is *g*_*c*_, *ρ*_*c*_, *a*_*c*_, *λ*_*c*_, *e*_*c*_, and *µ*_*c*_. As mentioned earlier, we consider metabolic and transport reactions (equations (1), (2), (7)) to be taking place on a faster time scale and the processes of protein expressions (equations (4),(5), (6), (8)) to be taking place at a slower time scale. For simplicity, we assume all fast processes are taking place at the same rate and slower ones are taking place at the same but with a low rate. In particular, we consider the faster processes are taking place at a rate that is 10 times that of the slower processes. We non-dimensionalize time with the rate of the slower dynamics of the system. All the results shown in the following sections are presented in these non-dimensionalized units.

## 3 Results and Discussion

We numerically solved the system equations (1) - (8) explicitly using numpy library in Python. For each case, the simulation was run until a steady state was reached. For the cases involving stochastic factors, each simulation was run multiple times to arrive at statistical convergence.

### 3.1 ATP metabolism in healthy articular cartilage

#### 3.1.1 Steady state response

We performed a numerical simulation of the model to obtain the steady-state response of the system in terms of ATP levels in AC. The model demonstrates that the ATP production in AC is via both modes of glycolysis and OXPHOS. The exact pathway depends on the availability of glucose and oxygen. As shown in Fig. 2A, at low glucose levels, the ATP production is primarily by glycolysis. An increase in the available glucose in the extracellular space (*g*_*o*_) results in a sudden transition of ATP synthesis pathway from glycolysis to OXPHOS (at *g*_*o*_ ≈ 0.3 in Fig. 2A). This transition from glycolysis to OXPHOS results in a nearly three fold increase in the ATP production for the parameter values studied here. This discontinuous transition from glycolysis to OXPHOS points towards bistability in the system. It shows that for low glucose levels glycolysis is the stable mechanism of ATP production. At high glucose levels, on the other hand, the system switches to the other stable fixed point, that is oxidative phosphorylation.

**Figure 2.**
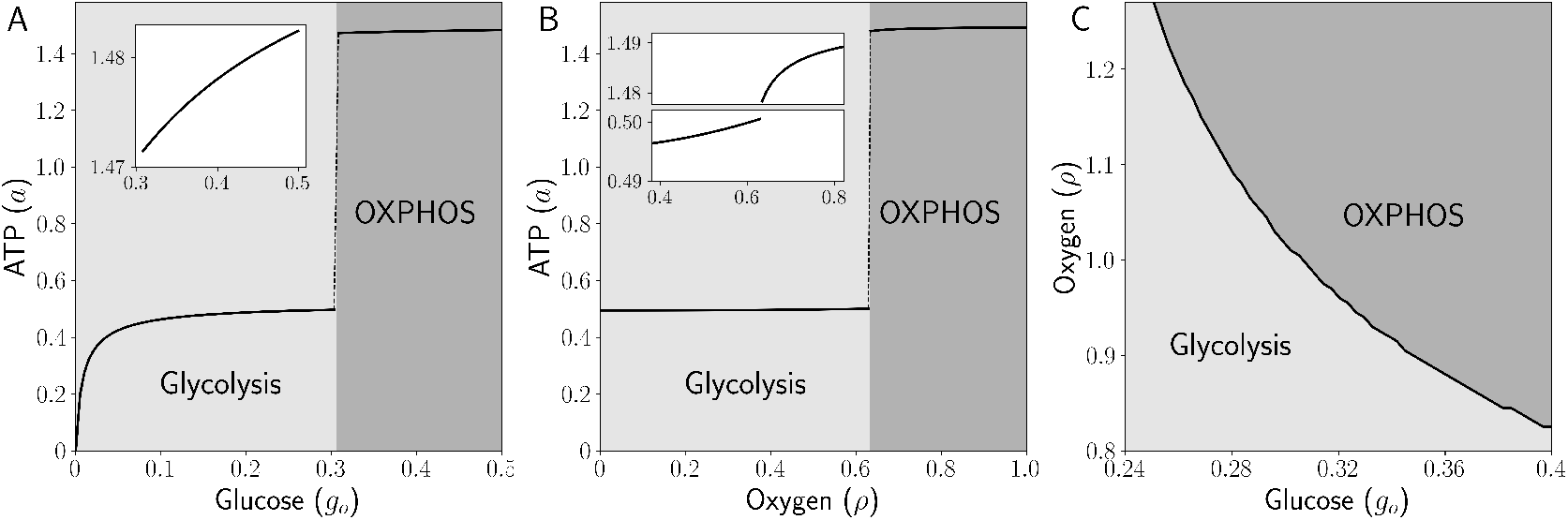
Steady state of ATP levels in healthy articular cartilage. Dependence of steady-state ATP levels in articular cartilage on (A) available glucose *g*_*o*_, and (B) oxygen *ρ*. At low levels of glucose (or oxygen) the ATP production is primarily via glycolysis. For larger glucose (or oxygen) levels it is produced by oxidative phosphorylation (OXPHOS). Insets in both (A) and (B) show enlarged versions of the respective curves near the transition point. (C) Phase diagram showing modes of ATP production in articular cartilage.

Similarly, oxygen availability also shows a similar bistable structure of the system with two fixed points (Fig. 2B). As expected, at low oxygen levels, the system follows glycolysis, an anerobic mechanism of ATP synthesis (Chaudhry and Varacallo, 2018). However, at high levels of available oxygen, it shifts towards OXPHOS, an aerobic mechanism of ATP synthesis. As shown in the inset in Fig. 2B, even though ATP levels increase with oxygen, its dependence on oxygen availability within each pathway is very weak. Fig. 2C shows the phase diagram mapping the dominant metabolic modes (glycolysis vs. OXPHOS) across physiological nutrient ranges. Notably, the boundary in the phase diagram underscores the synergistic dependence of OXPHOS on both glucose and oxygen availability, as neither substrate alone suffices to sustain mitochondrial ATP production.

The bistable behavior of oxygen availability highlights a critical aspect of cellular energy regulation in articular cartilage. Interestingly, within each metabolic state, ATP production remains relatively insensitive to fluctuations in oxygen concentration. This suggests a built-in metabolic robustness that protects cellular function against minor environmental changes. It further emphasizes that efficient mitochondrial ATP production requires both adequate glucose and oxygen, underscoring the need for a balanced nutrient environment to maintain cartilage health and prevent metabolic dysregulation.

### 3.1.2 Effect of stochastic perturbations

The extracellular space of articular cartilage is subject to stochastic fluctuations in glucose and oxygen availability due to multiple physiological factors. Dynamic mechanical loading during daily activities, such as walking or running, transiently alters nutrient diffusion from the synovial fluid into the cartilage matrix, creating variability in local glucose and oxygen concentrations (O’hara et al., 1990; Sampson et al., 2019). Additionally, inflammatory cytokines (not explicitly modeled in this study) can also disrupt metabolic demand by altering chondrocyte activity, while changes in vascular supply to adjacent tissues may indirectly perturb oxygen delivery (Goldring, 2000; Wojdasiewicz et al., 2014). The heterogeneous composition of the extracellular matrix further contributes to local noise by creating diffusion barriers that vary spatially, leading to uneven nutrient distribution. Together, these factors ensure that chondrocytes in vivo constantly experience stochastic perturbations in their metabolic microenvironment. We explored the effect of such stochastic noise in glucose and oxygen at the steady-state ATP levels in articular cartilage. We considered that the extracellular availability of glucose and oxygen are is given by

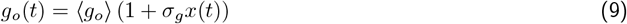

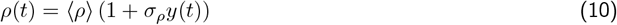

where *x*(*t*) ∼ N(0, 1) and *y*(*t*) ∼ N(0, 1) are random variables following normal distributions and *σ*_*i*_ (where *i* = *g, ρ*) represents the strengths of the stochastic noise. This form of stochastic noise is motivated by the fact that in biological systems fluctuations scale with mean concentrations, that is, higher baseline glucose can lead to larger absolute deviations (Ghosh et al., 2012). Furthermore, this multiplicative noise also prevents nonphysical negative concentrations, which is not the case for additive noise which can result in negative concentrations for ⟨*g*_*o*_⟩ *< σ*_*g*_ and ⟨*ρ*⟩ *< σ*_*ρ*_.

The effect of stochastic noise in the levels of glucose and oxygen on ATP levels in AC are shown in Fig. 3. It reveals a significant metabolic adaptation: oxidative phosphorylation emerges as the preferred ATP production pathway under fluctuating nutrient conditions. Specifically, we find that increasing the strength of stochastic noise in glucose (increasing values of *σ*_*g*_) and oxygen (increasing values of *σ*_*ρ*_)shifts the transition point between glycolysis and OXPHOS toward lower mean concentrations of both nutrients. This suggests that chondrocytes may rely more on OXPHOS than previously assumed, even in the relatively hypoxic environment of articular cartilage. Despite the strong effect on the transition point, the noise does not show any significant effect on the steady-state levels of average ATP in AC.

**Figure 3.**
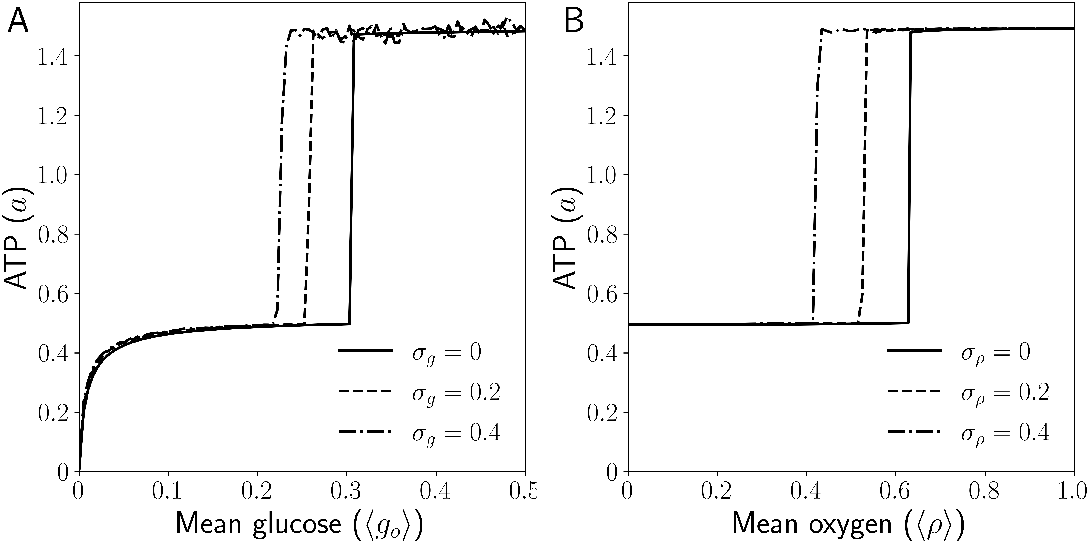
Effect of stochastic perturbations *σ*_*i*_ on steady state ATP levels. Steady state ATP levels in articular cartilage as a function of (A) mean glucose levels ⟨*g*_*o*_⟩ and (B) mean oxygen levels ⟨*ρ*⟩ in presence of stochastic noise.

### 3.2 Effect of acute hypoxia on ATP metabolism

As observed previously, the system shows bistable behavior, characterized by two stable steady states in ATP production. To investigate how acute hypoxia influences this metabolic balance, we simulated a transient oxygen deficit by reducing oxygen levels by a fixed amount Δ*ρ* for a finite duration *τ*, after which oxygen availability was restored to baseline (see dashed lines in Fig. 4A). This perturbation mimics physiologically relevant hypoxic episodes, such as those induced by mechanical loading or vascular fluctuations in articular cartilage (Imhof et al., 1997). We tracked the resulting ATP dynamics during this.

**Figure 4.**
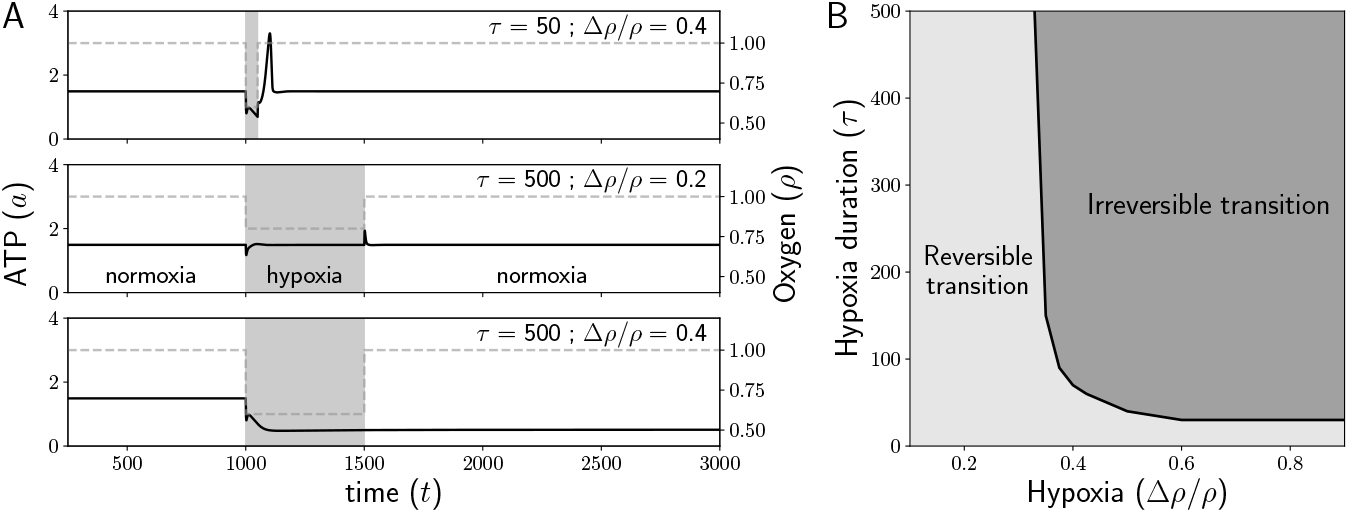
Hypoxia induced transition to glycolysis. (A) Time evolution of ATP levels in AC under the effect of acute hypoxia. For a smaller duration or a smaller drop in oxygen levels, ATP levels resume back to a normal level (top two panels). A larger drop in oxygen for a longer duration results in a transition to lower ATP levels (bottom panel). (B) Phase diagram illustrating the metabolic shift from oxidative phosphorylation to glycolytic ATP production following acute hypoxia.

#### 3.2.1 Hypoxia induced transition to glycolysis

Numerical simulations reveal that short-term hypoxia (small *τ*) or weak hypoxia (small Δ*ρ*) permit metabolic recovery once oxygen levels are brought back to normal (see top two panels in Fig. 4A). However, prolonged or severe oxygen deprivation drives the system toward glycolytic dependence (bottom panel in Fig. 4A). We also observe that once the metabolic shift takes place towards glycolysis, it does not change back to OXPHOS even if higher oxygen levels (higher than normal) or higher glucose levels (higher than normal) are made available (Fig. S1 in supporting information). This suggests that hypoxia induced transition from OXPHOS to glycolysis is irreversible through biochemical means. The phase diagram in Fig. 4B summarizes this metabolic shift from oxidative phosphorylation to glycolytic ATP production following acute hypoxia. We observe that the drop in oxygen levels needs to be sustained for a sufficient duration to induce a metabolic shift to glycolysis. For a large drop in oxygen, a shorter duration is sufficient to induce this transition, whereas a weak hypoxic episode might not result in the permanent metabolic shift.

These results reveal critical insights about the metabolic flexibility of chondrocytes, where recovery to OXPHOS upon reoxygenation post short hypoxia points to a resilience mechanism that aligns with physiological loading cycles. However, prolonged or severe hypoxia triggers an irreversible shift to glycolytic ATP production, persisting even after normoxia restoration, indicative of a biochemical hysteresis effect. This metabolic “lock-in” occurs only when hypoxia exceeds threshold combinations of duration and magnitude (Fig. 4B). The irreversibility of this transition suggests that chondrocytes lose metabolic plasticity under chronic hypoxia.

#### 3.2.2 Hypoxia-induced OXPHOS-glycolysis transition under stochastic perturbations

We have seen earlier that the presence of stochastic noise in the levels of extracellular glucose and oxygen affects the preferred metabolic pathway for ATP synthesis (Fig. 3). Thus, we investigated its role in the reversible OXPHOS-glycolysis switch during hypoxia-reoxygenation cycles.

Fig. 5 shows the critical boundary separating reversible and irreversible metabolic transitions in chondrocytes subjected to hypoxia-normoxia cycling. This analysis reveals that stochastic noise exerts a protective effect in both glucose and oxygen fluctuations, as increased variability consistently reduces the parameter space associated with irreversible glycolytic shifts. However, the magnitude of this protection differs substantially between nutrient types-while stochastic glucose variations (Fig. 5A) expand the recoverable OXPHOS region by a small margin, oxygen-level fluctuations (Fig. 5B) induce a pronounced shift in the transition boundary with increasing noise amplitude. This shows that the noise-dependent preservation of metabolic reversibility aligns with cartilage’s natural exposure to dynamic mechanical loading, where intermittent compression creates transient oxygen gradients.

**Figure 5.**
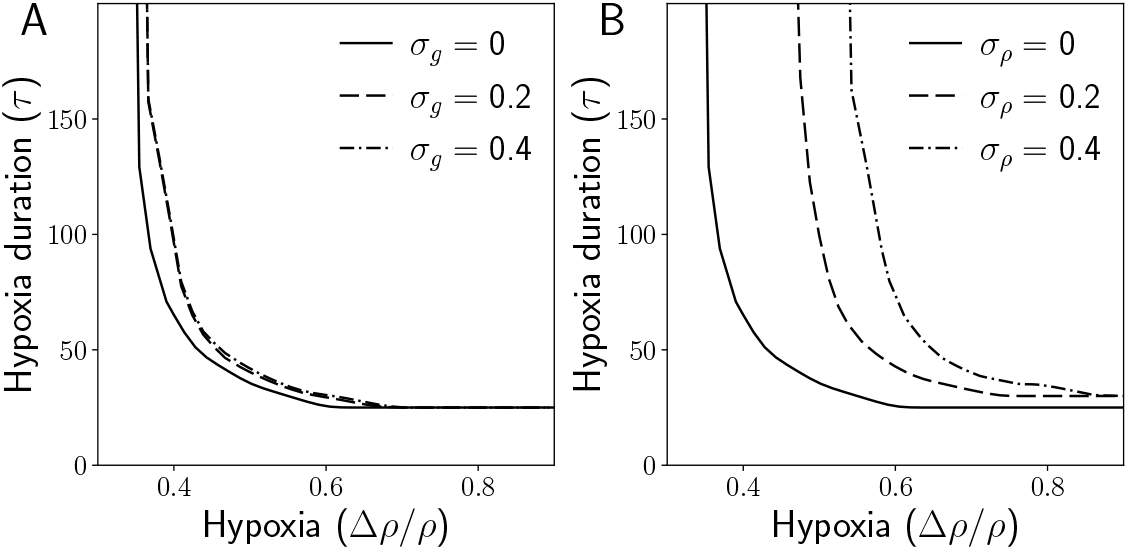
Effect of stochastic perturbations on OXPHOS-glycolysis transition. The boundary between reversible and irreversible transitions between oxidative phosphorylation and glycolysis for reference when stochastic noise is present in (A) glucose levels, and (B) oxygen concentration. Solid line in both panels is the same curve as that shown in Fig. 4B, that is, in the absence of any stochastic noise. Each curve divides the Δ*ρ/ρ* − *τ* parameter space into two regions: the lower-left zone, where OXPHOS recovers upon reoxygenation, and the remaining area, where the glycolytic shift becomes irreversible.

### 3.3 ATP metabolism and osteoarthritis

As mentioned earlier (Fig. 1B), it is known that sustained hypoxia-induced glycolytic metabolism in chondrocytes results in the production of advanced glycation end-products (AGEs) (Pi et al., 2024; Jiang et al., 2024; Saudek and Kay, 2003). These AGEs are known to disrupt the ECM levels by their enhanced degradation with the help of matrix metalloproteases (MMPs)(Saudek and Kay, 2003). Therefore, with the help of the model, we explored the role of hypoxia driven changes in ECM that have the potential to progress to osteoarthritis.

#### 3.3.1 Effect of hypoxia on ECM density

We numerically solved equations (7)-(8) with inputs from other equations. Here, we considered ECM dynamics under the hypoxic conditions described in Fig. 4. The time evolution of AGEs, ECM, and the levels of HIF-1*α* are shown in Fig. 6.

**Figure 6.**
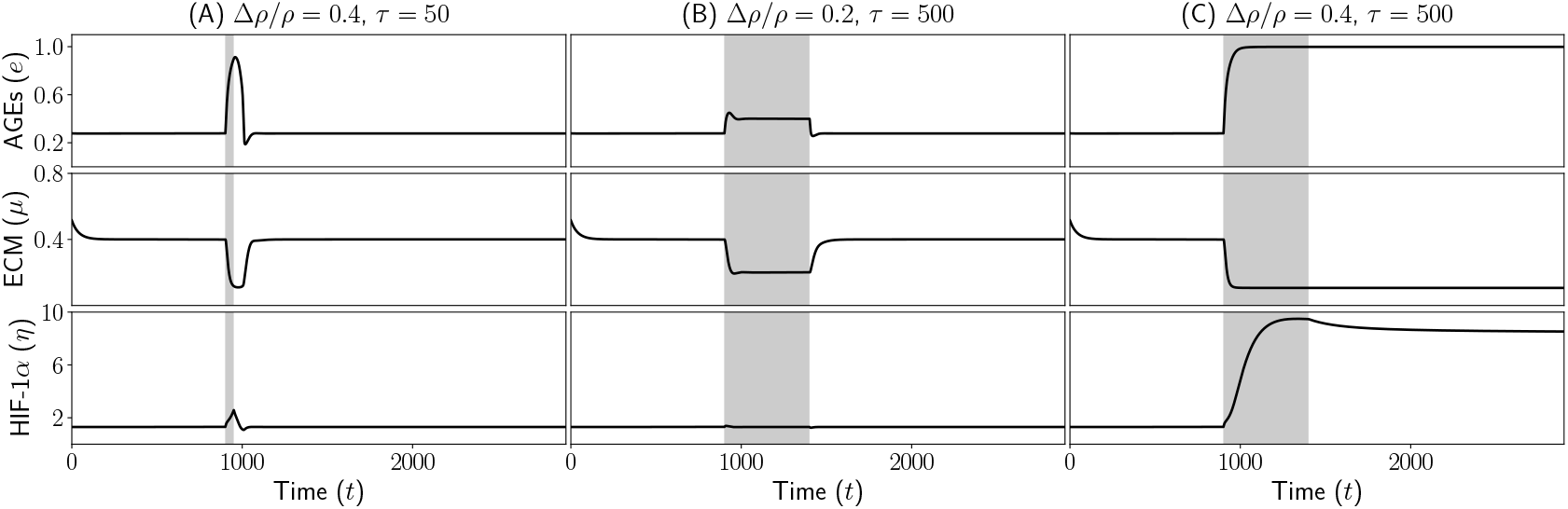
Effect of hypoxia on AGEs, ECM and HIF-1*α*. Plots show evolution of AGEs, ECM and HIF-1*α* under the effect of hypoxia of different magnitude and durations (same as in Fig. 4) - (A) Δ*ρ/ρ* = 0.4, *τ* = 50, (B) Δ*ρ/ρ* = 0.2, *τ* = 500, and (C) Δ*ρ/ρ* = 0.4, *τ* = 500. The period of hypoxia is shaded in gray.

As expected, a drop in levels of available oxygen results in an increase in the advanced glycation end-products (Fig. 6A-C). For shorter durations of hypoxia, it does not reach a steady state before normal oxygen supply is restored (Fig. 6A). In this scenario, the levels of all three entities revert back to their homeostatic levels (all three panels in Fig. 6A). We observe a similar pattern in the case of weaker hypoxia (Δ*ρ/ρ* = 0.2) for longer duration (*τ* = 500) where normal homeostatic levels are restored in normoxic conditions (Fig. 6B). For strong hypoxia (Δ*ρ/ρ* = 0.4) for a longer duration, on the other hand, results in a decrease in the ECM levels and an increase in AGEs and HIF-1*α*. These changes are not reversible under normoxic conditions (Fig. 6C). It needs to be noted that the irreversible decrease in the ECM levels is accompanied by an irreversible transition of ATP synthesis from OXPHOS to glycolysis (Fig. 4).

This metabolic reprogramming, driven by HIF-1*α* stabilization, suggests that chondrocytes under chronic hypoxia become trapped in a glycolytic state, impairing their ability to regenerate ECM even after oxygen levels return to normal. The irreversible ECM degradation aligns with OA pathology, where AGEs, with the help of MMPs, promote ECM degradation, resulting in reduced tissue resilience. These findings highlight hypoxia duration and intensity as key determinants of OA onset: transient hypoxia may be adaptive, but beyond a threshold, it triggers a pathological cascade of metabolic dysfunction, AGE-mediated ECM damage, and failed repair—hallmarks of early OA (Tonge et al., 2014; Yao et al., 2023).

Next, we tested for possible therapeutic strategies for the reversal of the osteoarthritic low density ECM state, which is triggered by the glycolytic switch induced HIF-1*α* stabilization, to a normal one. In the following, we explore four potential mechanisms for the reversal of the glycolytic switch that can help increase ECM density.

#### 3.3.2 Therapeutic potential of exogenous ECM on osteoarthritis

Exogenous ECM components, such as collagen hydrolysate and undenatured collagen, show promise as potential therapies for osteoarthritis, supported by preclinical and clinical studies (Honvo et al., 2020). Current evidence suggests that exogenous ECM supplementation could aid OA management, but more rigorous preclinical harmonization and large-scale clinical trials are required for definitive conclusions. We also tested this strategy using the model in the following manner. We consider the administration of exogenous ECM components by modifying equation (8) describing the dynamics of ECM as

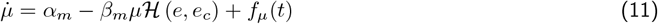

where the last term

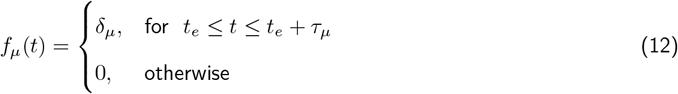

0, otherwise represents the exogenous ECM supplementation at a rate *δ*_*µ*_ for *τ*_*µ*_ duration after hypoxia period *t*_*e*_. Using this, we consider a long episode of hypoxia that results in ECM degradation (the pathological case shown in Fig. 6C) and estimate the ECM recovery with increasing magnitudes of *δ*_*µ*_. We assume that the exogenous ECM is supplied to the system right at the end of hypoxia for a duration *τ*_*µ*_.

As shown in Fig. 7, we find that the exogenous ECM supplementation on cartilage recovery under hypoxic conditions, a key factor in osteoarthritis progression, has dynamic effects. The data show that introducing exogenous ECM leads to a transient recovery of endogenous ECM levels, particularly noticeable for high values of *δ*_*µ*_. This recovery results in ECM levels rising significantly compared to hypoxia-only conditions. However, this improvement is short-lived; post-therapy, ECM concentrations gradually decline, eventually reverting to OA-like levels. This suggests that while exogenous ECM temporarily compensates for matrix loss, it fails to induce lasting restoration.

**Figure 7.**
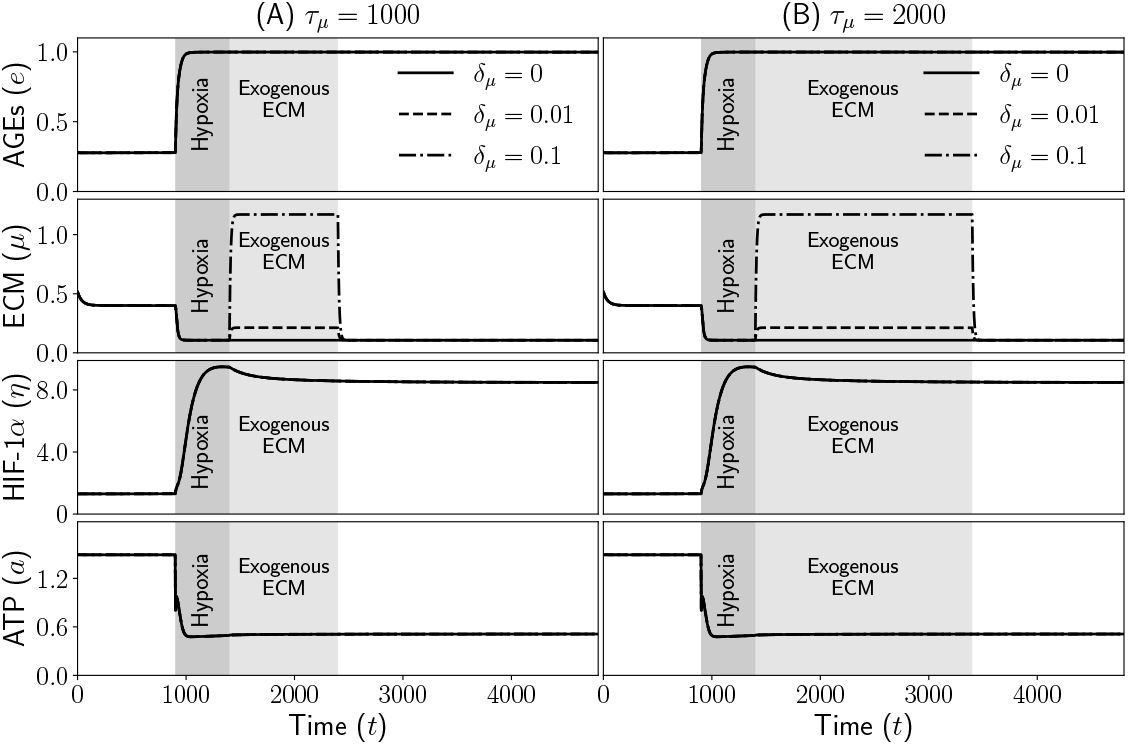
Exogenous ECM as potential therapeutic agent against ECM degradation. Time evolution of the levels of AGEs, ECM, HIF-1*α* and ATP after hypoxia with administration of exogenous ECM at different rates for two durations (A) *τ*_*µ*_ = 1000 and (B) *τ*_*µ*_ = 2000. The dark and light shaded regions mark the duration of hypoxia and administration of ECM components, respectively. For AGEs, HIF-1*α* and ATP the three curves in each panel coincide with each other.

The underlying mechanism appears tied to metabolic inflexibility in chondrocytes. Exogenous ECM supplementation does not alter the metabolic markers associated with hypoxia - specifically, HIF-1*α* and ATP levels remain unchanged compared to hypoxia-only groups. HIF-1*α*, a master regulator of hypoxic response, stays elevated, and ATP production remains suppressed, indicating a persistent glycolytic state. This metabolic stasis implies that exogenous ECM acts merely as a structural scaffold, unable to reprogram chondrocyte metabolism toward oxidative phosphorylation, which is critical for sustained ECM synthesis.

The transient efficacy of exogenous ECM highlights a key therapeutic limitation: without addressing the hypoxic metabolic shift, interventions relying solely on matrix replacement are palliative rather than curative. For long-term OA management, strategies must combine ECM supplementation with metabolic modulation (see next sections) to break the cycle of hypoxia-driven degradation.

#### 3.3.3 Stochastic perturbations induced reversal of glycolytic switch

As discussed earlier, stochastic fluctuations in glucose and oxygen levels can help sustain the oxidative phosphorylation pathway even when the average concentrations of these metabolites are low. Building on this observation, we investigated how random perturbations in glucose and oxygen availability influence cartilage recovery under hypoxic conditions. Time evolution of articular cartilage metabolism is shown in Fig. 8.

**Figure 8.**
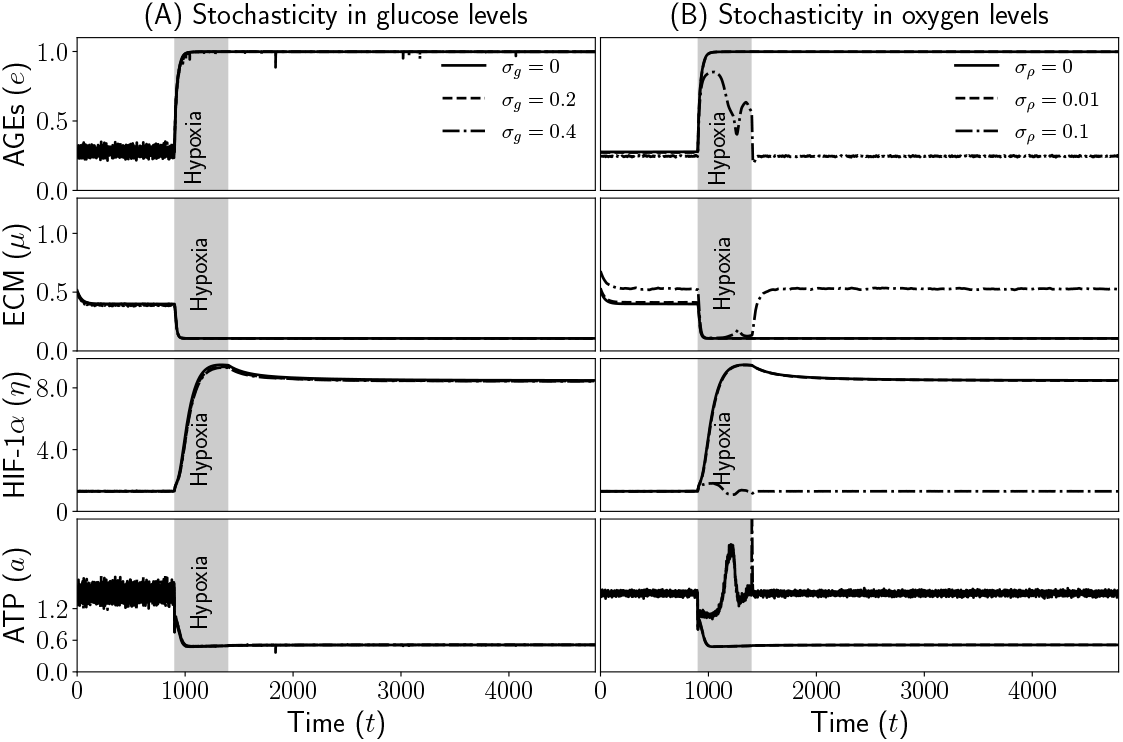
Effect of stochasticity in glucose and oxygen levels on glycolytic switch. Time evolution of the levels of AGEs, ECM, HIF-1*α*, and ATP after hypoxia in the presence of stochasticity in the levels of (A) glucose and (B) oxygen. The shaded regions mark the duration of hypoxia.

Fig. 8B reveals that introducing stochasticity in oxygen concentrations (*σ*_*ρ*_) promotes ECM recovery. However, stochasticity in glucose level did not show any remarkable effect on ECM recovery or ATP metabolism (Fig. 8A). Further, we also observe that higher noise amplitudes (*σ*_*ρ*_ = 0.1) in oxygen levels correlate with elevated baseline ECM levels compared to noiseless conditions (*σ*_*ρ*_ = 0). This suggests that physiological variability, such as intermittent supply of oxygen, may enhance ECM synthesis under steady-state conditions.

Despite this higher baseline ECM levels, during acute hypoxia, ECM levels drop precipitously across all stochasticity levels, converging to similar low values regardless of prior noise amplitude. This implies that while stochasticity bolsters baseline ECM production, it cannot fully counteract severe oxygen deprivation. Notably, post-hypoxic recovery is observed, with ECM levels rebounding to pre-hypoxia baselines, particularly in high-noise scenarios. This recovery phase highlights the resilience conferred by metabolic adaptability.

The findings bridge the mechanical and metabolic pathways of ECM regulation. While physical activity is traditionally linked to ECM upregulation via mechanotransduction (Leong et al., 2011; Zhao et al., 2020; Wang et al., 2023), our results suggest a parallel metabolic mechanism: stochastic oxygen variations - mimicking the dynamic microenvironment of the active joint - prime chondrocytes to resist and recover from hypoxic stress.

Higher stochasticity, likely reflecting increased physical activity, not only elevates baseline ECM but also ensures faster post-hypoxic recovery, possibly by maintaining metabolic flexibility (evidenced by ATP and HIF-1*α* dynamics). Thus, it underscores the dual role of physiological noise in cartilage homeostasis, advocating for therapies that harness natural metabolic variability (e.g., physical exercise based interventions) to delay OA progression.

#### 3.3.4 Enhanced ATP synthesis as a potential mechanism for OA reversal

As we have seen, ECM levels in articular cartilage and their regulation are closely linked with the metabolism, specifically ATP synthesis (Fig. 4). Motivated by this insight, we explored whether enhancing ATP production could serve as a mechanism to reverse the glycolytic switch and restore oxidative phosphorylation in normoxia. Specifically, we investigated how improvements in ATP availability influence the metabolic state transitions within the cartilage. For this, we modified the equation describing ATP dynamics as

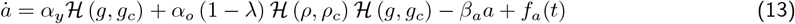

where the last term

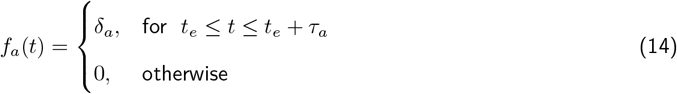

stands for enhanced ATP production lasting for *τ*_*a*_ duration after hypoxia ends at *t* = *t*_*e*_. The time evolution of the reversal dynamics is shown in Fig. 9.

**Figure 9.**
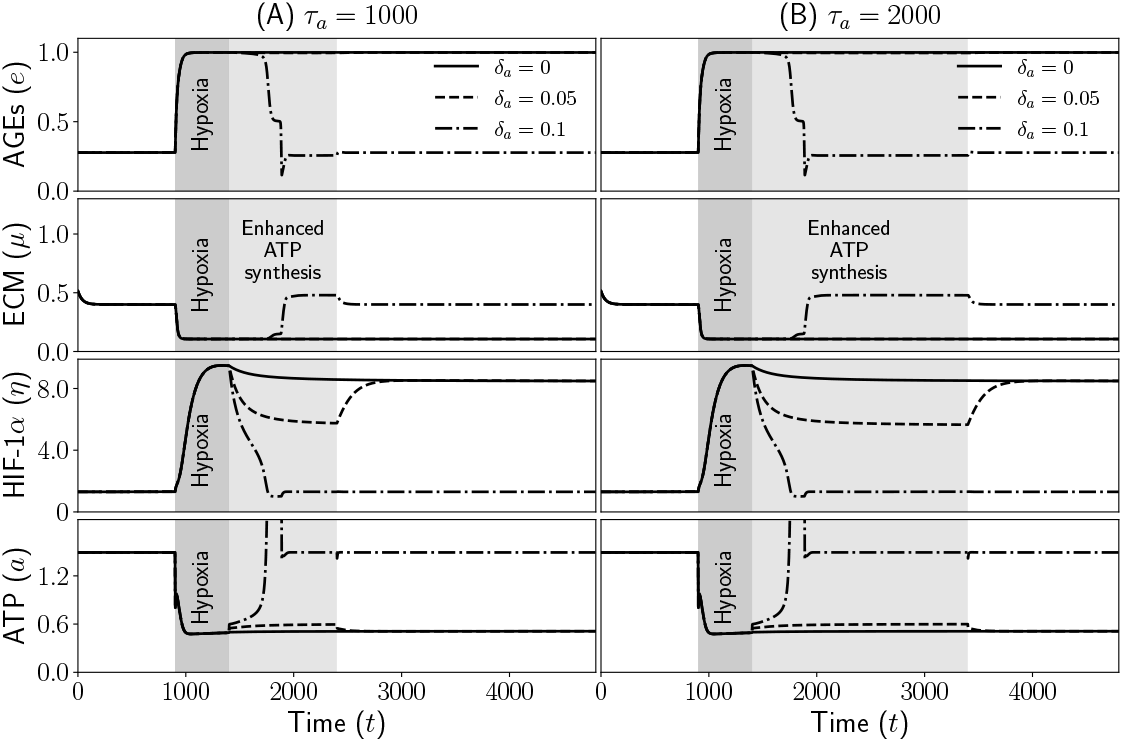
Effect of enhanced ATP synthesis in reversal of glycolytic switch. Time evolution of the levels of AGEs, ECM, HIF-1*α*, and ATP after hypoxia in the presence of enhanced ATP synthesis at different rates for two durations (A) *τ*_*a*_ = 1000 and (B) *τ*_*a*_ = 2000. The dark and light shaded regions mark the durations of hypoxia and enhanced ATP production, respectively.

We find that enhanced ATP production drives cartilage recovery following hypoxic stress, revealing a direct link between cellular energy metabolism and ECM restoration (Fig. 9). This recovery, however, is contingent on the magnitude of ATP elevation-higher ATP levels (e.g., *δ*_*a*_ = 0.1 in Fig. 9) correlate with more robust and sustained ECM restoration, particularly at longer durations (*τ*_*a*_ = 2000). Notably, we also find a mechanistic coupling between ATP and HIF-1*α*. Enhanced ATP synthesis coincides with reduced HIF-1*α* levels, suggesting that metabolic reprogramming away from glycolysis (a hallmark of hypoxia) suppresses HIF-1*α* stabilization. This inverse relationship underscores how ATP availability modulates the hypoxic response - elevated OXPHOS appears to “reset” chondrocyte metabolism, diminishing HIF-1*α* driven catabolic pathways that degrade ECM.

The transient nature of recovery in lower ATP-enhancement scenarios (*δ*_*a*_ = 0.05) implies a threshold effect: only sustained ATP elevation can durably counteract hypoxia-induced damage. These findings highlight ATP as a central regulator of cartilage resilience, proposing that therapies targeting mitochondrial function (e.g., OXPHOS boosters) could synergize with structural ECM therapies to halt osteoarthritis progression. These findings bridge metabolic and structural repair of ECM, emphasizing that energy homeostasis is necessary for long-term recovery from OA.

#### 3.3.5 HIF-1*α* as a potential target in osteoarthritis

Based on understanding the role of HIF-1*α* in articular cartilage metabolism, we explored whether targeted degradation or destabilization of HIF-1*α* could serve as a potential mechanism for reversing osteoarthritic changes. Specifically, we investigated how reducing HIF-1*α* levels influences ECM synthesis and cartilage recovery. To model this, we assumed the administration of a drug molecule that enhances the degradation of HIF-1*α*, applied over a fixed duration. This intervention modifies the dynamics of HIF-1*α* as

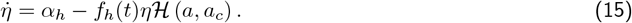

Where

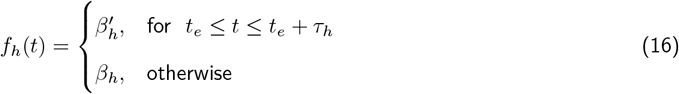

where 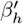 and *β*_*h*_ are the HIF-1*α* degradation rates with and without drug molecule, respectively, and 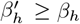. Using this setup, we analyzed how modulating HIF-1*α* degradation affects cartilage recovery under hypoxic conditions (Fig. 10).

**Figure 10.**
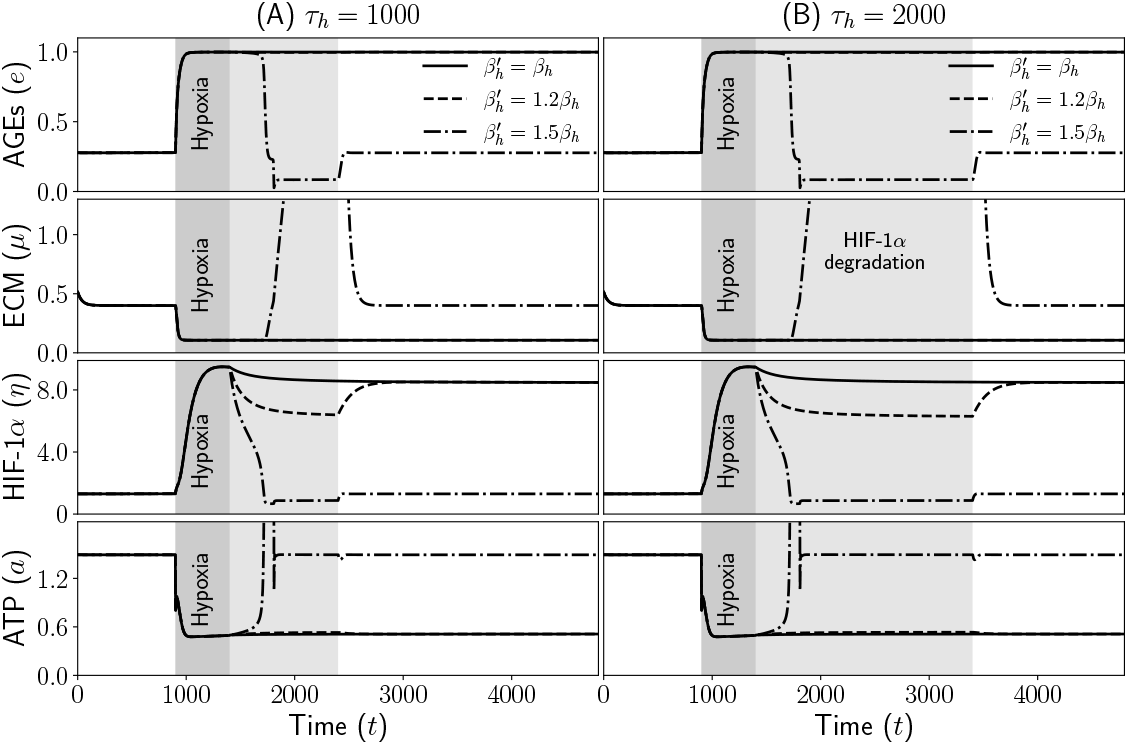
Effect of HIF-1*α* destabilization on glycolytic switch. Time evolution of the levels of AGEs, ECM, HIF-1*α*, and ATP after hypoxia with HIF-1*α* destabilization at different rates and durations. The dark and light shaded regions mark the durations of hypoxia and HIF-1*α* degradation, respectively. Due to low HIF-1*α* levels ECM concentrations go beyond the axis ranges.

Similar to ATP-driven recovery, enhanced HIF-1*α* degradation 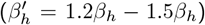 significantly improves ECM restoration post-hypoxia, demonstrating that HIF-1*α* suppression, whether direct or indirect, promotes ECM synthesis. However, unlike ATP modulation, HIF-1*α* degradation also leads to supra-physiological ECM accumulation (*µ* ≫ 1.0), which may stiffen articular cartilage and disrupt joint biomechanics, potentially worsening OA progression despite increased matrix content. This highlights the need for precise dosage optimization to balance ECM synthesis with tissue functionality. The dose-dependent effects of HIF-1*α* suppression further complicate therapeutic applications: higher degradation rates 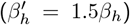 drastically reduce HIF-1*α* levels but may overshoot metabolic equilibrium, delaying ATP recovery, while moderate degradation 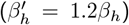 achieves a more harmonious restoration of both ATP and ECM levels. These findings suggest that partial HIF-1*α* inhibition, rather than complete ablation, may be optimal for maintaining chondrocyte homeostasis. This can be achieved through targeted HIF-1*α* modulation. For example, calibrated small-molecule inhibitors could replicate intermediate degradation rates to balance ECM synthesis and metabolic health (Rapisarda et al., 2002; Xia et al., 2009). Combining low-dose HIF-1*α* inhibitors with pro-glycolytic agents might sustain chondrocyte viability, while imaging techniques like MRI could monitor ECM stiffness in real time to prevent supraphysiological accumulation (Recht et al., 2001). Personalized medicine approaches (Chan and Ginsburg, 2011), including patient stratification based on baseline HIF-1*α* activity, could optimize dosage selection. Additionally, integrating HIF-1*α* suppression with mechanical loading regimens (resulting in stochastic perturbations in oxygen level, see Fig. 8), such as exercise, may help distribute ECM deposition more naturally and counteract cartilage stiffening.

This shows that, while HIF-1*α* degradation robustly reverses hypoxic damage, its potential to overdrive ECM synthesis requires careful clinical management. Future OA therapies must strike a balance between the role of HIF-1*α* in metabolism and matrix dynamics to restore joint function effectively.

### 3.4 Experimentally testable predictions and their physiological relevance

Now we summarize the main findings of the study from the perspective of experimental validation. These predictions offer insight into how metabolic regulation influences cartilage homeostasis and the progression or reversal of osteoarthritis. Each prediction suggests a potential therapeutic target that can be validated through specific experimental designs.

1. *Acute hypoxia triggers irreversible glycolytic switch*-The model has shown that exposure of chondrocytes to an acute hypoxic environment for long enough duration can result in irreversible switch to glycolytic metabolism (Fig. 4). Physiologically, this implies that even transient but severe ischemic events in the joints could cause lasting metabolic dysfunction, contributing to osteoarthritis progression. Therapeutic interventions must therefore not only restore oxygen delivery but also directly target metabolic pathways early after hypoxic insult to prevent irreversible shifts that degrade cartilage integrity.
2. *Enhanced ATP synthesis can reverse glycolytic shift*-We have also shown that one of the potential mechanisms of reversal of glycolytic shift is enhanced ATP synthesis (Fig. 9). This suggests that boosting mitochondrial ATP production could reverse harmful metabolic shifts, restoring cartilage function. Such an approach could offer a targeted therapy for early osteoarthritis, complementing existing treatments by stabilizing energy metabolism and promoting tissue repair at the cellular level rather than just managing symptoms.
3. *Stochastic oxygen fluctuations promote ECM recovery*-Another potential strategy for reversal of glycolytic switch and ECM restoration is an increase in the fluctuations in oxygen levels (Fig. 8). This can be brought about by physical activity, albeit in a controlled manner. This suggests that maintaining subtle oxygen variability could enhance cartilage resilience, offering a novel treatment strategy for osteoarthritis by promoting intrinsic repair mechanisms without invasive interventions.
4. *HIF-1α destabilization enhances ECM restoration*-HIF-1*α* has been a known therapeutic target in osteoarthritis (Zeng et al., 2022). The model presented in this work confirms this (Fig. 10). Physiologically, this suggests that fine-tuning HIF-1*α* activity post-injury could facilitate cartilage repair.

Together, these predictions provide a roadmap for bridging computational modeling with translational research in ATP metabolism and its role in osteoarthritis. They emphasize that regulating metabolic pathways can profoundly impact AC recovery and long-term joint health. Future experiments guided by these hypotheses could open new therapeutic avenues beyond traditional mechanical or anti-inflammatory treatments for osteoarthritis.

## 5 Conclusions

This study demonstrates that dysregulated ATP metabolism plays a central role in osteoarthritis progression by driving irreversible metabolic shifts in chondrocytes under chronic hypoxia. Our findings reveal that the hypoxia-induced glycolytic switch reduces ATP production and promotes ECM degradation through HIF-1*α* stabilization. While exogenous ECM provides temporary structural support, only interventions targeting ATP metabolism, either by enhancing OXPHOS or modulating HIF-1*α*, can reverse pathological changes. Importantly, physiological oxygen fluctuations from mechanical loading preserve metabolic flexibility, suggesting exercise may protect against OA by maintaining energy homeostasis. These results highlight ATP metabolism as a critical therapeutic target, emphasizing the need for combined metabolic and structural approaches to effectively treat osteoarthritis.

## Acknowledgements

We thank SERB and IITH for financial support.

